# ERASE: Extended Randomization for assessment of annotation enrichment in ASE datasets

**DOI:** 10.1101/600411

**Authors:** Karishma D’Sa, Regina H. Reynolds, Sebastian Guelfi, David Zhang, Sonia Garcia Ruiz, International Parkinson’s Disease Genomics Consortium (IPDGC), System Genomics of Parkinson’s Disease (SGPD), John Hardy, Sarah A. Gagliano Taliun, Kerrin S. Small, Mina Ryten, Juan A. Botía

## Abstract

Genome-wide association studies (GWAS) have identified thousands of genetic variants associated with various human phenotypes and many of these loci are thought to act at a molecular level by regulating gene expression. Detection of allele specific expression (ASE), namely preferential usage of an allele at a transcribed locus, is an increasingly important means of studying the genetic regulation of gene expression. However, there are currently a paucity of tools available to link ASE sites with GWAS risk loci. Existing integration methods first use ASE sites to infer cis-acting expression quantitative trait loci (eQTL) and then apply eQTL-based approaches. ERASE is a method that assesses the enrichment of risk loci amongst ASE sites directly. Furthermore, ERASE enables additional biological insights to be made through the addition of other SNP level annotations. ERASE is based on a randomization approach and controls for read depth, a significant confounder in ASE analyses. In this paper, we demonstrate that ERASE can efficiently detect the enrichment of eQTLs and risk loci within ASE data and that it remains sensitive even when used with underpowered GWAS datasets. Finally, using ERASE in combination with GWAS data for Parkinson’s disease and data on the splicing potential of individual SNPs, we provide evidence to suggest that risk loci for Parkinson’s disease are enriched amongst ASEs likely to affect splicing. Thus, we show that ERASE is an important new tool for the integration of ASE and GWAS data, capable of providing novel insights into the pathophysiology of complex diseases.

## Introduction

Over the last decade our understanding of complex genetic disorders has been transformed by the use of genome-wide association studies (GWAS) with 71,673 SNP-trait associations reported in the latest release of the NHGRI-EBI GWAS catalogue^1^. Complex neurological and neuropsychiatric disorders are no exception. The most recent GWAS for Parkinson’s disease (PD) reported 78 risk loci^2^, while that for schizophrenia^3^ reported 145. While there is a lag in terms of understanding the underlying biological processes involved in contributing to disease risk, it is thought that risk loci largely act by regulating gene expression as opposed to changing protein function^4^. This observation has led to an increasing interest in the genetic regulation of gene expression and the identification of expression quantitative trait loci (eQTLs, genetic variants that are associated with variation in gene expression) across the genome. eQTL studies have helped in the interpretation of GWAS findings across a range of disorders^5; 6^. However, one limitation of eQTL analyses is the requirement for large sample sizes. Since this can be difficult to achieve for some human tissues, such as specific brain regions, alternative approaches more robust to small sample sets are required.

Allele specific expression (ASE) analysis is another means of studying the genetics of gene expression, which is less dependent on sample size. This form of analysis quantifies expression variation between the two haplotypes of a diploid individual distinguished by heterozygous sites. While most heterozygous sites will have equal expression, this is not always true and differential expression at a transcribed position (termed an ASE site) can occur due to imprinting^7-10^, protein truncating variants triggering nonsense mediated decay^11; 12^ and cis-regulatory effects^13; 14^. ASE analysis being a within individual comparison, can be effective even with limited sample sizes as it is not confounded by factors such as age, sex or co-morbidity. Furthermore, since most ASE sites (particularly common SNPs) are driven by cis-regulatory effects^15-17^, ASE analyses could be integrated with GWAS to provide novel disease insights.

Existing tools for ASE-GWAS integration have been based on inferring cis-eQTLs from ASE data and then examining their association to disease^18; 19^. This approach requires an aggregation of information across the gene and can restrict the potential for inferring within-gene events relevant to the phenotype, namely splicing. Given that there is increasing evidence that common disease risk can be mediated through splicing^20-23^, this is a significant limitation. While there are other methods, which could be adapted for integration of ASE datasets with GWAS including SNPSnap^24; 25^ and stratified LD score regression^26^, they have been designed primarily with alternative forms of variant annotation in mind. For example, SNPSnap aims to provide control SNPs subsets for any set of SNPs. This implies accounting for a number of factors that become irrelevant when the source SNPs are ASEs (namely minor allele frequency and distance to the nearest gene, as ASEs are predominantly within exons). In stratified LD score regression^26^, ASE SNPs would be used as a form of annotation in heritability enrichment analyses. However, often ASE calling datasets are small, and thus would be underpowered in such an analysis. The intuitive approach would be to use ASE data directly with the GWAS using an ASE-specific method. However, there are intrinsic biases in ASE detection, which make such an approach challenging. ASEs are discovered only in transcribed regions, which means they are primarily located within exons, and are inevitably easier to detect confidently in a highly expressed genomic region. Given that GWAS loci are also enriched within regions as defined by open chromatin^27-29^ or tissue-specific expression features^3; 26; 30^, overlap between ASEs and GWAS hits have a high likelihood of occurring by chance. This makes it hard to know whether ASEs are truly providing useful disease insights.

In this paper we propose Extended Randomization for assessment of annotation enrichment in ASE datasets (ERASE), a tool that integrates ASE results with any SNP-based annotation dataset associated with a continuous score, such as eQTL or GWAS summary statistics. We test ERASE using multiple large publicly available ASE datasets. These datasets have been generated independently with different methods allowing us to test ERASE in the context of variable ASE calling pipelines. When we integrate the ASEs with an annotation dataset such as GWAS summary statistics, we take the shared set of SNPs, namely the SNPs both analysed for their ASE potential and their effect on the phenotype of interest. We then ask if ASE sites are enriched for loci linked to disease more than we would expect by chance. We use a randomization-based approach to assess enrichment. This approach has proven useful in the context of eQTLs^24^. ERASE has an added advantage in that it does not require individual-level genotype data to test whether detected ASEs are enriched for loci with higher scores in the annotation datasets and to determine the contribution of each annotation dataset to the enrichment. When ERASE compares the observed ASE calling set with a randomly chosen set of non-ASE SNPs it matches for the read depth. This is an important step because it ensures that any enrichment seen is due to the presence of an ASE and does not simply reflect high expression of a given gene. Moreover, we further extend this method to incorporate multiple annotation datasets in our enrichment analyses in order to better understand the specific properties of the ASEs. We do so by studying the relative contribution of each annotation dataset to the enrichment.

## Material and Methods

### ERASE

In this section, we describe the specifics of the method. We also provide a publicly available R package, available from GitHub (see Web Resources), implementing it.

#### ERASE - single annotation enrichment (SAE)

We wish to test whether significant ASEs in a study are also enriched for notable results in a SNP-based annotation dataset and apply a version of parametric bootstrapping to address this problem (Figure 1). Let the values in the annotation dataset be transformed such that large positive values represent more significant results (for example, a minus-log_10_ transformation would achieve this for p-values). Let *A* be the set of SNPs tested for ASE, let *G* be the set of SNPs in the SNP annotation dataset and let *C* be the common SNPs present in both *A* and *G*. Let *Ase(*+) be the set of SNPs in *C* with a significant ASE signal, and let *Ase(*−) be the set of SNPs in *C* with a non-significant ASE signal. We use *Ase(*−) to model the null distribution of SNPs across the genome. To control for the confounding effect of read depth, SNPs are randomly selected with replacement from *Ase(*−) to create a SNP set with the same size and read distribution as that of *Ase(*+). This is done by placing the SNPs in *Ase(*−) into bins based on their average read depth across samples (using read depth information from the study that generated the ASE data). By default, a bin corresponds to an interval of average reads with width 2, such that SNPs with average read depth of 3 and 4 will be in one bin while those with a read depth of 5 and 6 will be in the next bin and so on. All SNPs with an average read depth greater than 200 are placed in a single final bin. For each SNP in the *Ase(*+) set, a ‘null’ counterpart is selected at random from the corresponding bin that matches its read depth, until a full set of ‘null’ SNPs is obtained. By default, the random selection of ‘null’ SNP sets is repeated 10^4^ times, and the mean annotation SNP score (for example, mean(−log_10_ (p-value)) if a GWAS summary statistic dataset is used) is calculated for each set.

**Figure 1.**
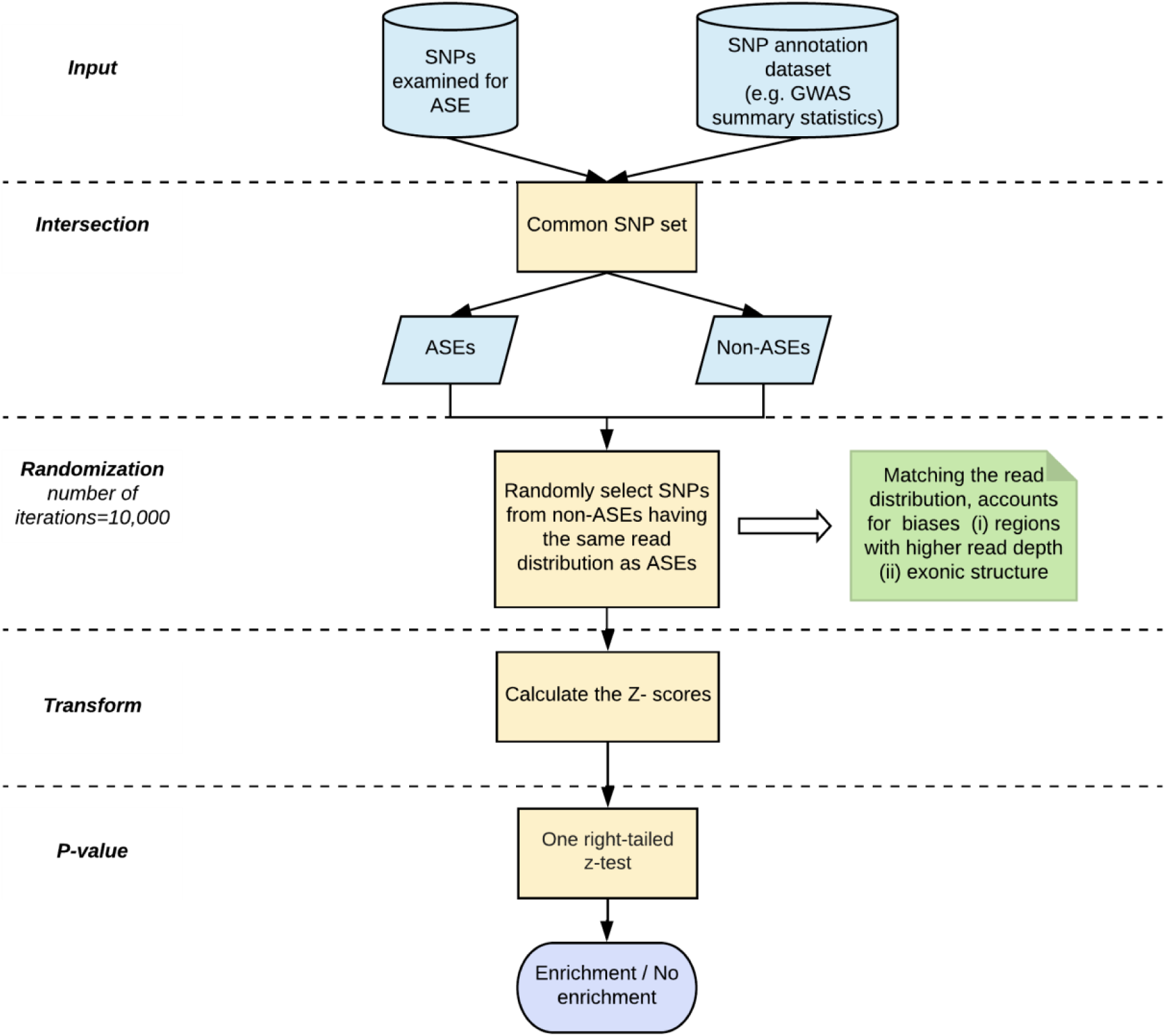
Overview of ERASE-single annotation enrichment (SAE) method. The figure represents a graphical overview of the general approach used in ERASE. The ERASE-SAE version of the tool just needs as input the set of SNPs examined for ASE and the SNP annotation dataset. It starts with finding the SNPs common (C) to both datasets. To construct a null distribution of random samples, the non-ASE SNPs from C are randomly selected 10^4^ times such that their average read distribution matches that of the ASE SNPs in C. This accounts for biases in ASE discovery due to high read depth and exonic structure. The mean and the standard distribution obtained from the randomization are used to calculate the z-score for the observed set of ASEs. The p-value is generated by a one right-tailed z-test.

Let *r*_*i*_ be the mean annotation SNP score for the *i*-th iteration. The mean 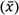 and standard deviation *σ*(*x*) of the random *r*_*i*_ values are used to calculate the z-score for the observed mean value from our set of significant ASE SNPs, 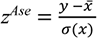, where y is the mean SNP score for the *Ase(*+) SNP set. The final p-value for the enrichment of SNPs for the annotation amongst ASEs versus non-ASEs is obtained from a one right-tailed z-test on *z*^*Ase*^.

#### ERASE - multiple annotation enrichment (MAE)

The multiple annotation version of ERASE (Figure S1) tests simultaneously for enrichment of ASEs in more than one SNP annotation dataset. Simultaneous enrichment in two or more SNP annotation datasets means that we effectively test whether the same ASE SNPs tagging SNPs in one source also tag SNPs in the other, by using only SNPs common to the ASE calling and SNP annotation datasets. Starting from this basic concept, we also examine the relative contribution of each annotation dataset to the enrichment. To illustrate the approach in the case of two annotation datasets, we will consider one set of SNPs, *G*, with GWAS summary association statistics, and another set of SNPs, *S*, with transcript specificity values (as obtained, for example, from the SPIDEX^31^ database). As with ERASE-SAE, we assume the annotation values are expressed in a way that large positive values represent more notable results.

To perform the ERASE-MAE test, we will consider the set of SNPs, *C*, which are common to *A, G* and *S*. The randomization procedure proceeds in a similar way as for ERASE-SAE, but is now run twice, once for each of the two annotation types. Let *Ase(*+) be the set of SNPs in *C* with a significant ASE signal, and let *Ase(*−) be the set of SNPs in *C* with a non-significant ASE signal. For each annotation type, subsets of SNPs are randomly selected from the *Ase(*−) set using the same matching procedure as in ERASE-SAE, and the mean value of the relevant annotation scores are stored. By default, the randomization step is repeated 10^4^ times. In order to provide a consistent basis for comparison between the two annotation types, the stored values are then standardized by subtracting their respective means and dividing by their respective standard deviations, to create two sets of transformed values, 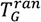 and 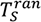. Similarly, the observed mean annotation values in the *Ase(*+) set are standardized in the same way to create two further transformed values, 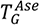 and 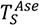.

The scores for the two annotations sources are then integrated as follows. Consider, *α* ∈ *[0,1]* to be a calibration parameter representing the relative weight assigned to each SNP annotation dataset, i.e. *α* being the weight for *G* and (*1* − *α*) the weight for *S*. We define the observed value of significant ASE sites in the integration of *G* and *S* as

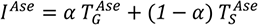

and the integrated value for the *i*th random set is then

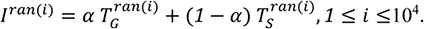

The p-value for the enrichment of *G* and *S* SNPs amongst ASEs versus non-ASEs is obtained with a one right-tailed z-test using the z-score for *Ase*, calculated using the mean and standard deviation of the *I*^*ran*(*i*)^, 1 ≤ *i* ≤ 10^4^ values and the *I*^*Ase*^ for a given value of *α*. We assume that the population values of 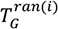 and 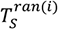 are both normal (i.e. z-scores measure the deviation from the mean, in standard deviation units), thus *I*^*ran*(*i*)^ should also be normal. Note that the value of *I*^*Ase*^depends upon the relative contribution of each SNP annotation dataset. Therefore, we would aim to find the value of *α* that optimizes *I*^*Ase*^(i.e. the one that yields the highest enrichment). The extension of the method for the N annotation datasets case follows naturally when we define a series of weights *w*_*i*_ for N, such that 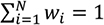. Finding the value of the weights when N=2 is reduced to vary one of the weights (i.e. alpha) in [0,1] such that

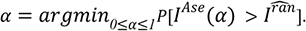

When N ≥ *3*, the brute force approach we suggest using (i.e. trying a representative set of value combinations for all *w*_*i*_’s) is less applicable, due to computational complexity. Approaches to deal with this problem in a computationally efficient manner exist. For example, using dynamic programming we can decompose finding the N weights problem into two subproblems: finding the values for N-1 weights corresponding to dimensions 1…N-1 and the value for *w*_*N*_in the N-th dimension. And again, we can decompose finding the N-1 weights into finding N-2 weights and weight *w*_*N*−1_. Once we reach directly solvable problems, we can reconstruct composite solutions backtracking towards the initial problem. We are currently exploring this approach.

### Stratified LD score regression (LDSC)

Stratified LDSC^26^ partitions common SNP-heritability (estimated from GWAS summary statistics) across customizable sets of SNPs; in this case we use ASE annotations. All annotations were constructed in a binary format (i.e. using 1 if the SNP was present within the annotation and 0 otherwise), using only SNPs with a minor allele frequency > 5%. Annotations were added individually to the baseline model of 53 annotations provided by Finucane et al. (version 1.1, see Web Resources), comprising genome-wide annotations reflecting genetic architecture, such as coding regions and histone marks. HapMap Project Phase 3 (HapMap3)^32^ SNPs and 1000 Genomes Project^33^ Phase 3 European population SNPs were used for the regression and LD reference panels, respectively. The MHC region was excluded from all analyses due to the complex and long-range LD patterns in this region. For all stratified LDSC analyses, in addition to the enrichment, enrichment p-value and regression coefficient, we report a one-tailed p-value (coefficient p-value) based on the coefficient *z*-score outputted by stratified LDSC. A one-tailed test was used as we were only interested in annotation categories with a significantly positive contribution to trait heritability, conditional upon the baseline model.

## Results

### Datasets

ERASE’s ability to handle relatively low sample numbers emerges from the fact that ASE analyses work well with small sample sizes. Thus, the best suited dataset for ASE analyses is one based on specific brain regions, which can be hard to obtain in large numbers. With this in mind, we tested ERASE on ASE calling datasets generated from human brain and lymphoblastoid cell lines. The former was generated from 49 putamen and 35 substantia nigra tissue samples, of the UK Brain Expression Consortium dataset^34^. These samples were obtained from neuropathologically normal individuals of European descent. All sites having > 5 reads in a sample were tested for ASE. In some analyses, data from putamen and substantia nigra were considered together, to distinguish between lack of power and no real enrichment, and the minimum FDR was taken across both tissues. The second ASE calling dataset, comprised of lymphoblastoid cell lines (LCLs)^15^ from 462 individuals of the 1000 Genomes Project where sites with at least 30 reads and both alleles seen were tested for ASE. From both the ASE calling datasets, SNPs with an FDR < 5% were considered ASE sites.

We used 1000 Genomes SNP data^35^ (phase 3 genotype data – see Web Resources) to simulate a set of genome-wide SNPs representing null ASEs. The 1000 Genomes data was selected as it has formed the basis of analyses in many studies^26; 36; 37^. SNPs were selected from 503 individuals of European ancestry from the following 5 populations: Utah Residents (CEPH) with Northern and Western European Ancestry (CEU), Toscani in Italia (TSI), Finnish in Finland (FIN), British in England and Scotland (GBR), Iberian Population in Spain (IBS). The 3 types of SNP annotation datasets we used to test ERASE are eQTLs, GWAS full summary statistic datasets and the alternative splicing prediction SPIDEX database. We used the gene-level eQTL data generated from 111 putamen and 69 substantia nigra tissue samples of the UK Brain Expression Consortium dataset^34^ to test enrichment of eQTLs amongst ASEs. Depending on the ASE dataset analyzed, different GWAS summary datasets were used when testing for enrichment of disease risk loci. For UKBEC-putamen dataset, Schizophrenia^3; 38; 39^, PD (excluding 23andMe data)^40^ (access obtained, with permissions from IPDGC and SGPD), waist-to-hip ratio adjusted for body mass index (BMI)^41^ were used, with the former two used as positive controls and the latter as a negative control. In the case of the LCL dataset, rheumatoid arthritis^42^ and epilepsy^43^ were used as positive and negative controls respectively. GWAS summary data files for individuals of European ancestry were downloaded (see Web Resources) for consistency with the origin of individuals in the ASE calling dataset. Waist-to-hip ratio adjusted for BMI summary statistics were converted from hg18 to hg19 using the liftOver tool^44^. The other summary statistics already used hg19 co-ordinates. SPIDEX^31^ database (see Web Resources) serves as data source for transcript specificity while examining the specific properties of ASEs. It is a database of pre-computed list splicing variants with the percent inclusion ratio of the exon where the variant is present (ΔΨ). It reports the maximum ΔΨ value across 16 tissues.

### ERASE detects known enrichment of eQTLs amongst ASEs

Besides imprinting and nonsense mediated decay, it is known that cis-regulation can also generate ASE signals^13^. In fact, this process has been previously shown to account for most of the ASE sites^15-17^. We tested whether ERASE detects the known enrichment of eQTLs amongst ASEs using ASEs and gene-level eQTLs identified in human post-mortem putamen^34^. We applied ERASE-SAE to the 154,214 SNPs, which had been analysed for both evidence of allelic imbalance and eQTL activity. ASEs were placed into bins, based on the average read depth overlying the SNP of interest. We did the same for SNPs negative for the ASE test (termed non-ASEs). They were placed into bins for random selection such that the average read depth of the randomly selected SNPs showed a similar distribution as that of the ASEs. As expected, we found that the ASEs were enriched for gene-level eQTLs in putamen (1.96 × 10^−150^, Figure S2).

We repeated this analysis varying bin width (from 2 to 10) to model the impact of less accurately matching the read distribution of non-ASEs to ASEs and also calculated enrichment when no attempt was made to control for differences in read depth. We observed that with increasing bin sizes the p-value for eQTL enrichment amongst ASEs is stable across the bins but when read depth was totally ignored as a confounding factor, a highly inflated p-value was observed (Figure S2, File S1). Enrichment examined with a negative control, waist-to-hip ratio adjusted for BMI^41^ GWAS summary dataset, resulted in a significant p-value (2.23 × 10^−2^) being observed only when read depth was not accounted for. This suggests that controlling for read depth is effective and can help prevent a false positive result. Moreover, in this analysis we note the following trade-off: while the smallest bin width of 2 may provide us with the most accurate estimate of enrichment, since the average read distribution of the randomly selected SNPs closely matches that of the ASEs it is also true that the proportion of unique SNPs randomly selected on each iteration is lower than when larger bin sizes are used (Figure S3, File S2). This is simply because the pool of SNPs which are non-ASEs shrinks with the bin size. Thus, the choice of average read intervals needs to be balanced against the ability to generate independent randomizations. The higher the number of non-ASE SNPs with respect to ASEs falling within each bin, the more independent randomizations we will obtain. Having said this, we note that enrichment p-values were reasonably stable across a range of average read depth intervals in our test case (Figure S2).

### ERASE yields a controlled Type I error rate

We conducted simulation studies to estimate the Type I (false positives) error rates incurred by ERASE using the example of eQTL enrichment amongst ASEs detected in putamen^34^. This analysis was performed by generating null ASE datasets by selecting subsets of SNPs from amongst those identified within the 1000 Genomes dataset. To ensure consistency with the origins of individuals from the UKBEC dataset, only SNPs from individuals of European ancestry in the 1000 Genomes dataset were considered (see Datasets section for details). The generation of the null ASE datasets was performed accounting for the main biases affecting ASE discovery. SNPs were further selected on the basis of biallelic status, exonic location and presence within a gene known to be expressed in human putamen^34^. This led to the identification of 595,348 SNPs that could be considered as false ASEs, and these were used to construct null ASE calling datasets. Each time we tested enrichment of gene-level eQTLs against the null, we randomly selected 8,527 SNPs (matching the number of true ASEs that overlapped with the putamen gene-level eQTL summary SNPs) and ran ERASE with the gene-level eQTL dataset. In order to get enough null experiments, we repeated the process 1000 times and the Type I error rate was estimated using the proportion of ERASE tests yielding enrichment (ERASE p-values < 0.05). We found an acceptable ratio of Type I errors of 4.3%. (Figure 2, File S3).

**Figure 2.**
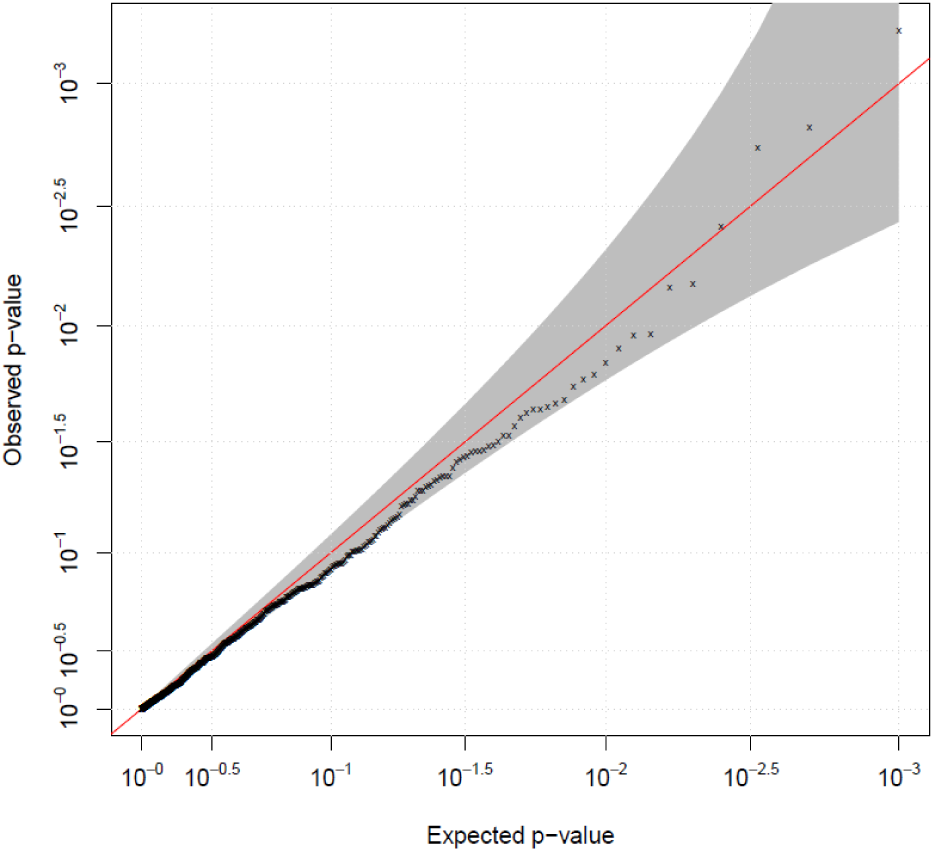
QQ-plot of simulations performed to assess Type I error. The estimate for the Type I error in ERASE we report is obtained by using the tool 1000 times with random sets of SNPs as ASEs. In each simulation, we generate null ASE datasets from the 1000 Genomes SNPs and test them for enrichment with ERASE. The tested enrichment is on gene level eQTLs from human post-mortem putamen samples. The figure shows a QQ-plot representing the null population of p-values we obtain. The grey area region marks the not significant p-values region. The pool of SNPs selected from the 1000 Genomes for the simulations were from individuals of European ancestry, of biallelic status, exonic in location and present within a gene known to be expressed in human putamen. The Type I error rate estimate (i.e. proportion of p-values greater than 0.05 out of the 1000 null p-values) obtained in the simulation was 4.3%.

### ERASE detects known enrichment of GWAS risk loci for rheumatoid arthritis amongst ASEs identified in lymphoblastoid cell lines

Given that ERASE was developed to enable the integration of publicly available ASE data with disease-relevant annotation datasets, such as GWAS data we wanted to test our method in this context. With this in mind, we investigated the enrichment of risk loci for rheumatoid arthritis and epilepsy, selected to serve as a positive and negative controls, amongst ASEs identified in lymphoblastoid cell lines (LCLs)^15^, a commonly used model of B-lymphocytes^45^ with regulatory sites already implicated in a range of autoimmune diseases. ERASE-SAE was run separately on the 51,403 and 35,199 sites that were examined for ASE and present in the rheumatoid arthritis^42^ and epilepsy^43^ GWAS summary datasets respectively. A bin size of 10 was used to enable a better sampling of the non-ASEs such that they matched the average read distribution of the ASEs in the LCL dataset. In line with previous studies, we found that the ASEs in LCLs were enriched for rick loci linked to rheumatoid arthritis (p-value = 1.40 ×10^−79^) and not for epilepsy (p-value = 0.19). Thus, we demonstrate the utility of ERASE by successfully running it on two independent, public ASE datasets.

### ERASE detects significant enrichment of risk loci amongst ASEs in underpowered GWAS’s

Studies have shown that GWAS’s require large sample sizes to achieve adequate statistical power across a full range of risk allele frequencies and effect sizes^46; 47^. As a consequence, for many diseases, including schizophrenia, GWAS analyses have been repeated using increasing numbers of cases and controls^3; 38; 39; 48^ with the aim of finding more risk loci each time. We tested ERASE at varying levels of GWAS size in order to calibrate its behavior in potentially underpowered conditions. This analysis was performed by running ERASE on three schizophrenia GWAS summary datasets with varying sample sizes (including cases and controls) ranging from 33,100 (Ripke et al., 2013^38^) to 105,318 (Pardiñas et al., 2018^3^) (Table 1). We examined if ERASE could detect enrichment of risk loci for schizophrenia in putamen ASE signals when sample sizes were small, noting that medium spiny neurons of the putamen (amongst other cell types including pyramidal cells and interneurons) have recently been implicated in the pathogenesis of schizophrenia^49^. As ASEs generated tend to be in highly expressed genes and these tend to be neuronal in the case of schizophrenia, not accounting for read depth as a confounder can result in high expression being considered as the reason for any enrichment observed. Controlling for read depth ensures that the enrichment seen is due to the ASEs. We tested for enrichment of loci linked to waist-to-hip ratio adjusted for BMI^41^ as a negative control. As a comparison to our method, we used stratified LD score regression^10^ (LDSC) (see Material and Methods), a recently developed and popular tool. As expected, ERASE detected an enrichment of schizophrenia risk loci amongst ASEs identified in putamen using the largest available schizophrenia dataset^3^ (40,675 cases, 64,643 controls) (Table 1) with no significant enrichment for waist-to-hip ratio adjusted for BMI (p-value = 0.13). Similarly, stratified LDSC demonstrated a significant enrichment of schizophrenia heritability amongst ASEs (coefficient p-value = 1.82 × 10^−2^) with no significant enrichment for waist-to-hip ratio (coefficient p-value = 0.07). However, while ERASE was able to detect the enrichment of risk loci amongst ASEs even when schizophrenia GWAS datasets were smaller and consequently less well-powered, the enrichment of heritability amongst ASEs was non-significant when stratified LDSC was used with smaller schizophrenia GWAS datasets due to the larger error margins. Thus, we demonstrate that ERASE performs favorably in terms of detecting enrichment of risk loci in GWAS with small sample sizes.

**Table 1.** Enrichment of risk loci for schizophrenia in putamen ASEs observed with ERASE and LDSC, using schizophrenia GWAS’s of varying power. * indicates p-values and co-efficient p-values < 0.05.

| Schizophrenia GWAS | Number of cases | Number of controls | ERASE p-value | Stratified LDSC coefficient p-value |
| --- | --- | --- | --- | --- |
| Ripke et al.<sup>38</sup> | 14,395 | 18,705 | *6.01x10<sup>-12</sup> | 1.23x10<sup>-1</sup> |
| Schizophrenia Working Group of the Psychiatric Genomics Consortium<sup>39</sup> | 33,640 | 43,456 | *2.90x10<sup>-21</sup> | 6.55x10<sup>-2</sup> |
| Pardiñas et al.<sup>3</sup> | 40,675 | 64,643 | *1.52x10<sup>-20</sup> | *1.83x10<sup>-2</sup> |

### ERASE detects existing enrichment when ASEs are under-called

Next, we tested the sensitivity of ERASE in detecting the enrichment of risk loci amongst small ASE calling datasets (i.e. varying levels of ASE test stringency). We approached this by down sampling true ASEs. This analysis was performed by calculating the enrichment of risk loci for schizophrenia as determined by Pardinas and colleagues^3^ amongst decreasing percentages of ASEs identified in human putamen and substantia nigra^34^. In order to ensure this process was robust, for each subset of ASEs generated (including 5% and 10% increments-5%, 10%, …, 30%, 40%…, 90%) the random subsampling was repeated twenty times for each tested set, varying the random seed. We applied both ERASE-SAE and stratified LDSC to these ASE datasets. As expected, both ERASE and stratified LDSC generated a positive correlation between ASE dataset size and the significance of enrichment observed. However, using ERASE a significant enrichment of schizophrenia risk loci amongst ASEs was observed (median p-value = 4.01×10^−11^; Bonferroni cut-off = 2.07 × 10^−4^), even when the ASE dataset was down sampled to 30%, equating to 4,543 SNPs (Figure 3(a), File S4). On the other hand, stratified LDSC did not identify a significant enrichment of heritability for schizophrenia even when 80% of the original ASE annotation was included (median coefficient p-value at 80% = 5.27 × 10^−4^; Bonferroni cut-off = 4.95×10^−4^) (Figure 3(b), File S5). It only manages to reach a significant enrichment with all the ASEs (coefficient p-value = 4.19×10^−4^). We noted that in the case of stratified LDSC, enrichment in heritability within the ASE annotation was always observed but as the ASE annotation increased in size the confidence in these estimates increased, enabling identification of a significant coefficient p-value (Figure S4). Thus, we see that ERASE identifies a significant enrichment of risk loci within smaller subsets of ASEs compared to stratified LDSC.

**Figure 3.**
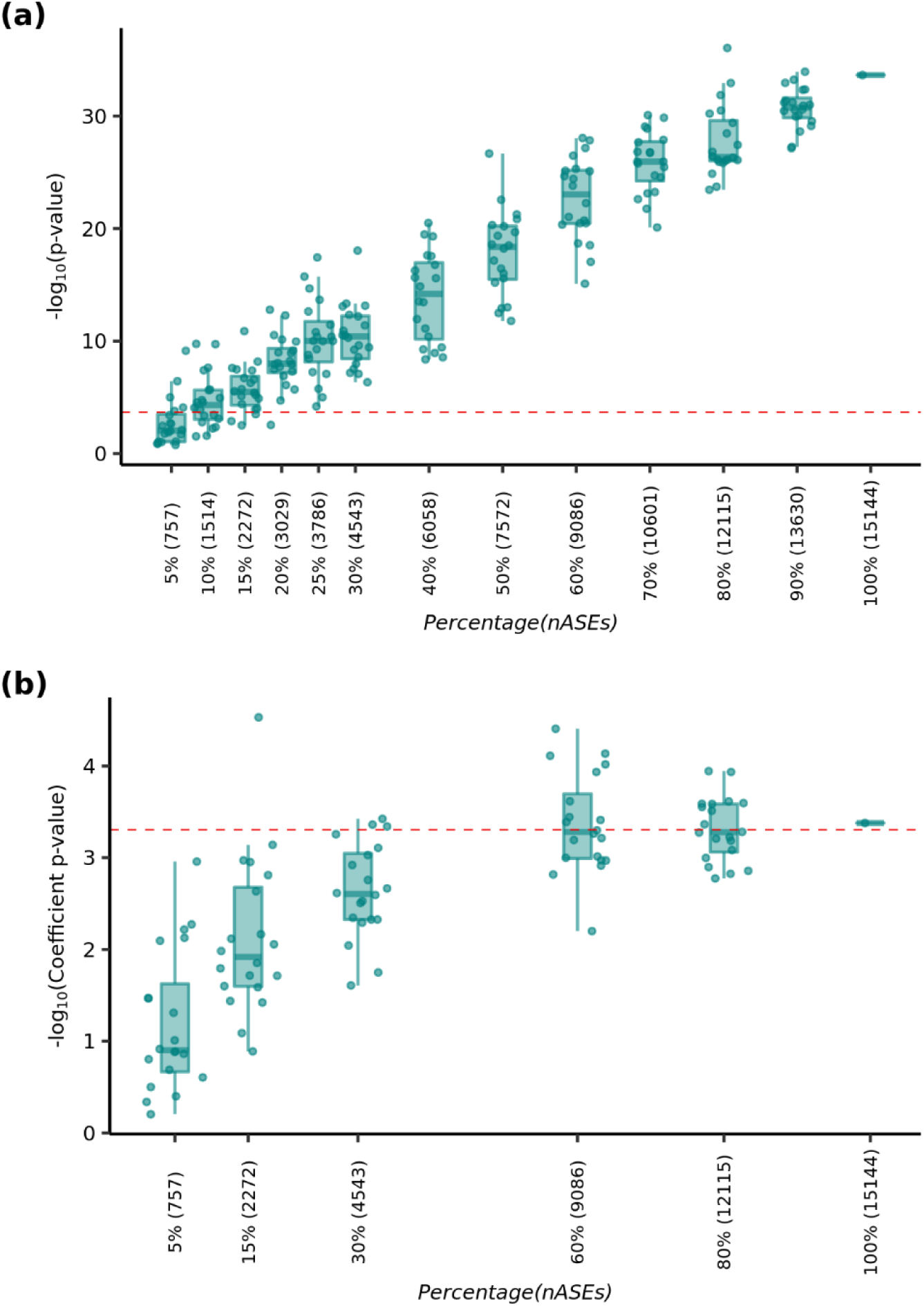
Assessment of sensitivity in ERASE and stratified LDSC with ASE SNP datasets of various sizes, when testing for observed enrichment of schizophrenia risk loci (Pardiñas et al., 2018). ASEs derived from putamen and substantia nigra were randomly subsampled to form sample sets of various sizes and 20 sample sets per size varying the random seed. These sample sets were tested with ERASE and stratified LDSC. Each box plot corresponds to the p-values or coefficient p-values, obtained with (a) ERASE or (b) stratified LDSC respectively, tested on the 20 sample sets. The x-axis shows the different sampling sizes of ASEs, i.e. the percentage of ASEs sampled from the total, followed by the number of ASEs that it equates to, within brackets. The y-axis shows the −log_10_ scale for the p-values or coefficient p-values. The red dashed lines indicate the corresponding Bonferroni cut-off applied to the results obtained from each of the tools. (a) ERASE identifies significant enrichment of schizophrenia risk loci amongst the ASEs (total of 15,144 signals) even when the number of ASEs was lowered to 30% (4,543 signals) of the total ASEs. (Bonferroni cut-off = 2.07 × 10^−4^). (b) Stratified LDSC identifies significant enrichment of heritability for schizophrenia only when the entire ASE annotation (15,144 signals) was included (Bonferroni cut-off = 4.95×10^−4^). Notably, an enrichment in heritability within the ASE annotation was always observed with stratified LDSC and as the ASE annotation increased in size the confidence in these estimates increased, enabling a significant coefficient p-value to be identified (Figure S4).

### ERASE demonstrates ASEs predicted to have stronger effects on splicing, drive enrichment of risk loci

ERASE works well when testing the enrichment of ASE in single annotation datasets, as evidenced by its ability to detect enrichments of schizophrenia and PD GWAS loci in ASEs called in putamen and substantia nigra^34^. ERASE also allows the integration of more than one annotation datasets, thus providing further insight into the nature of these ASE signals. To illustrate this, we used ERASE-MAE to examine if ASEs derived from putamen and substantia nigra, which were predicted by Guelfi et al. to have stronger splicing effects^34^ also drive the enrichment of risk loci linked to schizophrenia^3^ and PD^40^. We used SPIDEX^31^, a database of splicing variants, generated using machine learning, with the percent inclusion ratio of the exon where the variant is present (ΔΨ), for transcript specificity.

ERASE-MAE was applied to the set of 30,892 SNPs that were common to the ASE calling dataset and two annotation datasets, namely schizophrenia GWAS summary data and the SPIDEX variant dataset. Similarly, we applied this approach to the 35,922 SNPs common to the ASE calling dataset, PD GWAS summary data and the SPIDEX variant dataset. In each case, enrichment was examined by varying the relative importance of the two annotation datasets (the GWAS and SPIDEX) when calculating enrichment amongst ASEs. This was done by assigning the calibration parameter, α, which indicates the relative weight of the GWAS, with values ranging from 0 to 1 with 0.1 increments, such that an α value of 1 indicates an enrichment p-value obtained by using the GWAS annotation dataset alone, while an α value of 0 refers to an enrichment p-value based on the SPIDEX annotation dataset alone. We observed a very significant enrichment of risk loci for schizophrenia and PD amongst ASEs, as predicted by the previously observed enrichment of schizophrenia and PD risk loci amongst ASEs derived from putamen and substantia nigra (Figure 4, File S6).

**Figure 4.**
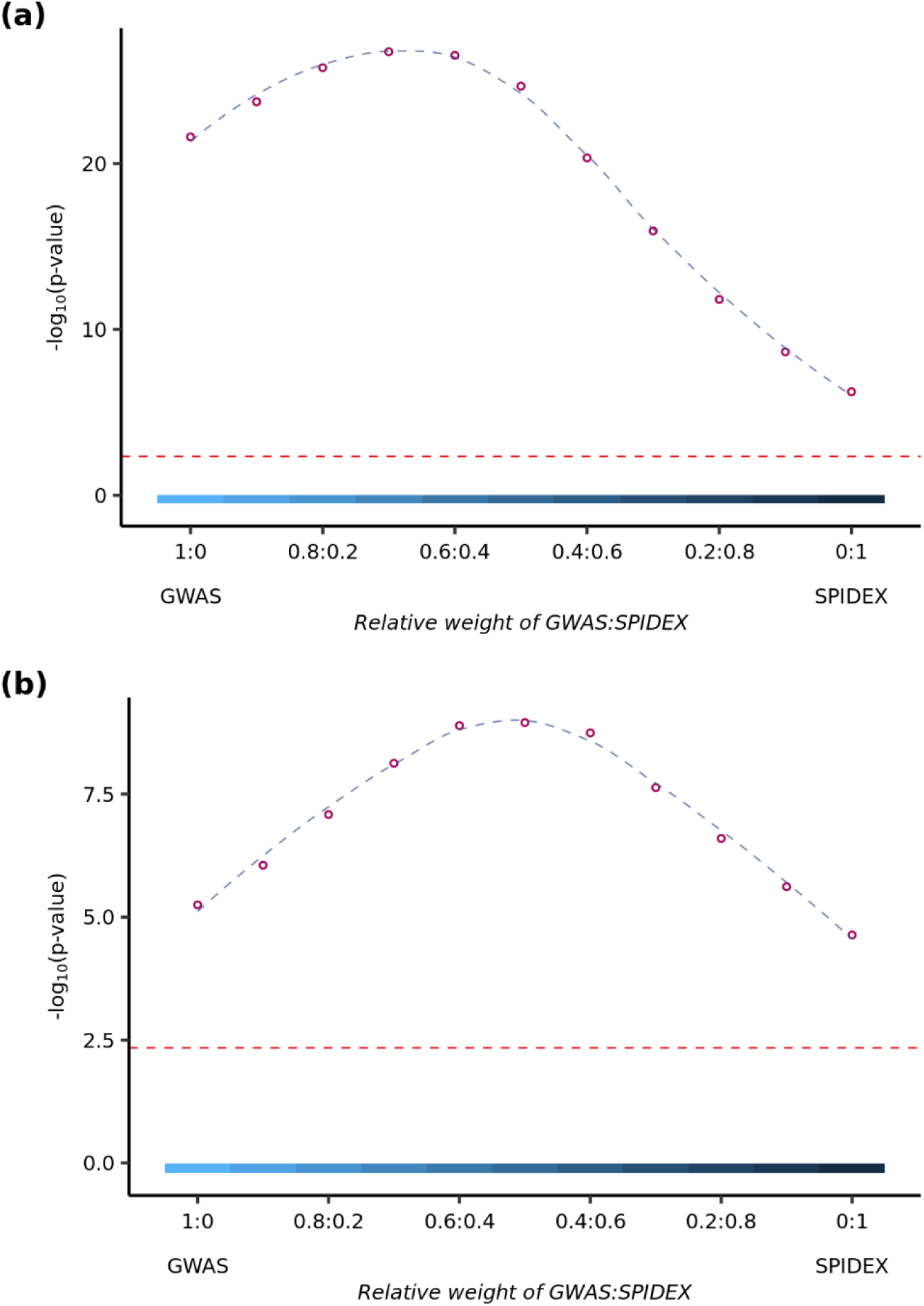
Results of ERASE-MAE, with varying the relative contributions of the two SNP annotation datasets to the final enrichment. ERASE-MAE was applied to ASEs derived from human putamen and substantia nigra using SNP annotation datasets SPIDEX and (a) Schizophrenia (Pardiñas et al. 2018), (b) Parkinson’s disease (Nalls et al.2018). The relative contribution of each SNP annotation dataset to the final enrichment was varied by assigning α, a calibration parameter, a range of values from 0 to 1 incremented by 0.1, such that an α value of 1 indicates the enrichment contribution is from the GWAS dataset alone and value of 0 from SPIDEX. The x-axis shows the relative weights assigned to GWAS and SPIDEX. The y-axis shows the −log_10_ scale for the p-values. The red dashed line indicates the Bonferroni cut-off of 4.55×10^−3^. In plot (a) we see an enrichment peak for schizophrenia risk loci and transcript specificity at 07:03. This reflects clearly that both contribute to the enrichment. It also suggests that the contribution is unbalanced, i.e. the contribution of schizophrenia risk loci is more relevant. In plot (b) the case is different. We see a balanced case, 0.5:0.5, in which both PD risk loci and SPIDEX equally contribute.

In addition, using ERASE-MAE and integrating the SPIDEX annotation, further increased the enrichment of risk loci amongst ASEs for both diseases. For schizophrenia, an enrichment peak was observed when the relative weight of GWAS to SPIDEX was 0.7:0.3 (p-value = 1.80 × 10^−27^) (Figure 4(a)), while for PD an enrichment peak was observed when the relative weight of GWAS to SPIDEX was 0.5:0.5 (p-value = 1.11 × 10^−9^) (Figure 4(b)). Thus, factoring in the splicing potential of ASEs improved the enrichment of risk loci for schizophrenia and PD. This suggests that the ASEs in these tissues are likely to be disease-associated splice sites.

## Discussion

ASE is a potentially powerful tool due to its ability to provide regulatory information even in the context of small numbers of RNA sequencing samples. However, the effectiveness of ASE analyses in understanding the biology of complex genetic diseases has been limited by the challenges of robustly integrating ASE data with other forms of annotation. This is analogous to existing methods used to calculate GWAS enrichment amongst eQTLs. In this study, we address this issue by presenting a method, ERASE, to calculate enrichment of a ranked annotation dataset amongst a set of ASEs. Using ASE data from two different public datasets and integrating this with cis-eQTL and GWAS data, we show that ERASE can reliably enable this form of analysis with a low Type I error rate (4.3%). We also demonstrate that read depth is a confounder when examining enrichment in ASEs and not accounting for it results in a highly inflated p-value. It should be noted that a key component of ERASE is its ability to control for read depth, an important bias in ASE detection. This is achieved by randomly sampling non-ASE sites within bins, such that they have a read depth distribution similar to that of ASEs. While the default bin size (set at 2) provides a conservative enrichment p-value, the proportion of unique SNPs randomly selected may be low when the numbers of non-ASE sites at the matching read depth are limited. This means that the bin size selection needs to be balanced against the proportion of unique SNPs. However, we also show that the enrichment p-value is not inconsistent or highly inflated at varying bin sizes, the key point being read depth is accounted for.

ERASE is easy to use requiring only the SNP set examined for ASE, with their corresponding read depths and whether an ASE was observed or not, and a ranked SNP annotation dataset. By directly using ASEs rather than leveraging ASE data across the gene to infer cis-acting eQTLs, ERASE makes use of all available ASE data and in particular retains ASEs, which are generated through splicing QTLs. This is important given the growing recognition that risk loci for complex diseases can operate by affecting transcript usage^21; 31; 50; 51^ and may explain the sensitivity of this method in the context of smaller GWAS or ASE analyses. ERASE detected the enrichment of schizophrenia risk loci amongst ASEs identified in human putamen across all schizophrenia GWAS analyses tested including where the smallest GWAS was used. While this enrichment was validated using an independent approach, namely stratified LDSC, this was only possible using the larger schizophrenia GWAS. Similarly, we found that even when the ASE sites identified in basal ganglia were down-sampled to only 4,543 SNPs (30% of all ASE sites) ERASE was still able to detect a significant enrichment of schizophrenia risk loci amongst ASEs. In the case of stratified LDSC, while enrichment in heritability was still apparent amongst this subset of ASE sites, the enrichment was not significant. Given the growing interest in using iPSC-derived cell types to identify cell and state-specific regulatory sites^52-54^, the sensitivity of ERASE is particularly relevant as generating, differentiating and maintaining the large iPSC-derived sample sets required for eQTL analysis remains time-consuming and expensive, while applying ASE analyses to small iPSC-derived sample sets is more feasible.

Importantly, we show how ERASE, in particular ERASE-MAE, can be used to provide insights into the pathogenesis of complex diseases. From a methodological standpoint, ERASE-MAE is useful to test ASE enrichment in more than one source simultaneously, namely whether the same subset of ASE SNPs enriches in all annotation sources. However, it can also help unravel unequal contributions from different annotation sources to the global enrichment. We illustrate this utility by showing that the enrichment of schizophrenia and PD risk loci is driven in part by the splicing-related properties of ASE sites. Our results provide additional evidence for the role of splicing in schizophrenia and PD, where rick loci have already been found to be enriched for splicing QTLs identified in human frontal cortex^22; 55; 56^.In summary, ERASE, a sensitive and robust tool, has been designed to use ASE data directly, assessing if it is enriched for SNPs in single or multiple annotation datasets. ERASE-MAE is the arm of the tool that evaluates the enrichment in more than one SNP annotation dataset and the relative contribution of each annotation to the overall enrichment. Notably, ERASE uses all ASEs rather than summarizing information across a gene, thus retaining ASEs generated due to splicing QTLs. Given the growing popularity of ASE analyses ^57-60^ and the increased interest in splicing, ERASE’s ability to detect enrichment in small ASE datasets and multiple annotations has the potential to add significantly to biological knowledge.

## Supplemental Data

Supplemental Data include four figures and 6 tables (as excel files).

## Acknowledgments

Mina Ryten, Karishma D’Sa, David Zhang and Sonia Garcia Ruiz were supported by the UK Medical Research Council (MRC) through the award of Tenure-track Clinician Scientist Fellowship to Mina Ryten (MR/N008324/1). Regina Reynolds was supported through the award of a Leonard Wolfson Doctoral Training Fellowship in Neurodegeneration. Sebastian Guelfi was supported by Alzheimer’s Research UK through the award of a PhD Fellowship (ARUK-PhD2014-16). This work was supported by the UK Dementia Research Institute. Full consortia acknowledgements are available in the supplemental data file (Supplemental Text 2)

## Web Resources

1000 Genomes genotype data, Phase 3, was downloaded on 18/4/18 from ftp://ftp.1000genomes.ebi.ac.uk/vol1/ftp/release/20130502/; Baseline LDSC annotations, https://data.broadinstitute.org/alkesgroup/LDSCORE/; LDSC, https://github.com/bulik/ldsc; SPIDEX, was downloaded on 27/2/17 from https://www.deepgenomics.com/spidex-noncommercial-download; UK Brain Expression Consortium data http://braineacv2.inf.um.es/; GWAS summary datasets downloaded were Schizophrenia ^3; 38; 39^ from https://www.med.unc.edu/pgc/results-and-downloads, Waist-to-hip ratio adjusted for body mass index (BMI)^41^ from the GIANT consortium website https://www.broadinstitute.org/collaboration/giant/index.php/GIANT_consortium_data_files, Rheumatoid arthritis^42^ from http://plaza.umin.ac.jp/~yokada/datasource/software.htm, Epilepsy^43^ from http://www.epigad.org/gwas_ilae2014/; ERASE https://github.com/karishdsa/ERASE

