## Supplemental figures for "ERASE: Extended Randomization for assessment of annotation enrichment in ASE datasets"

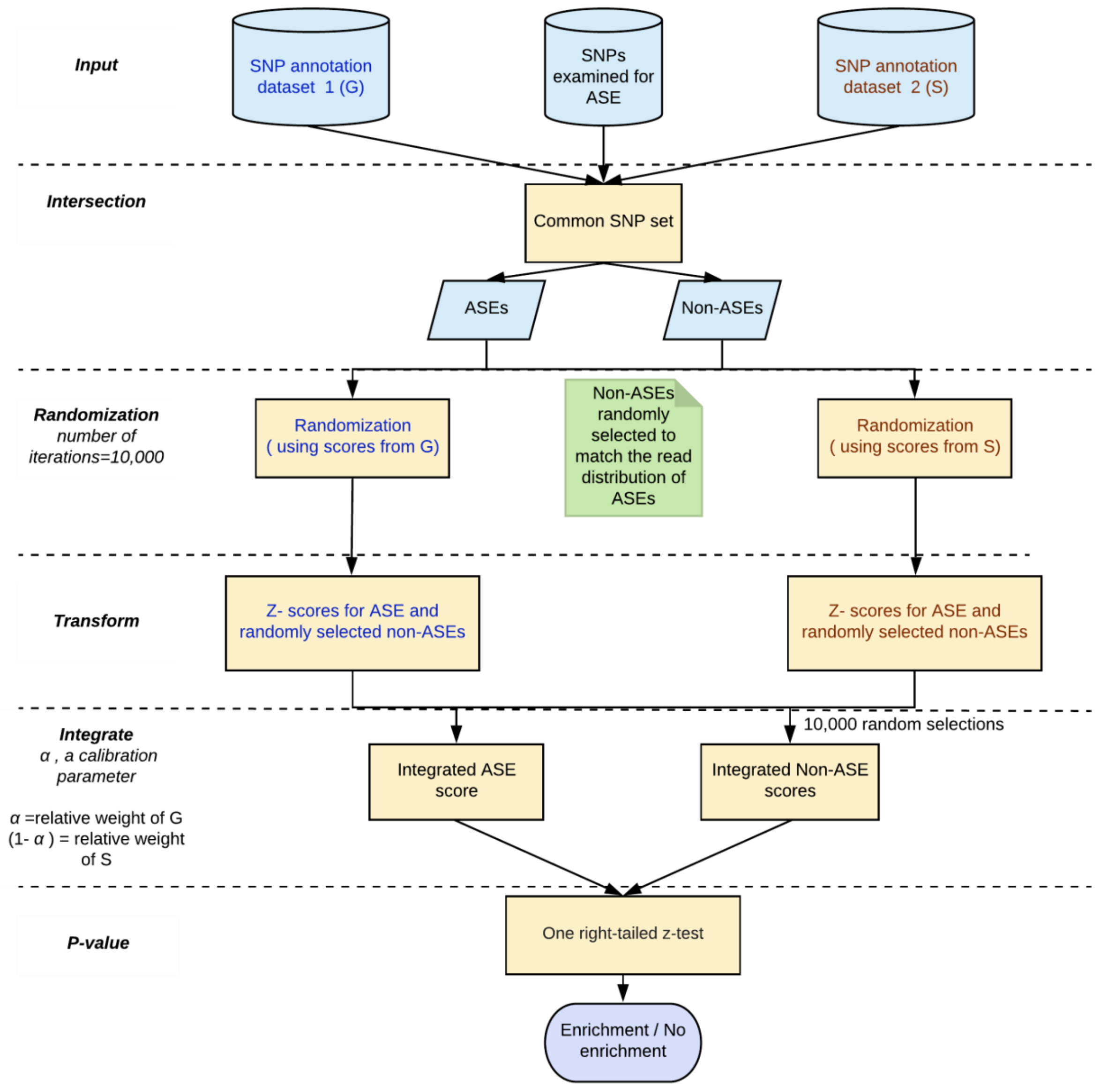


**Figure S1: Overview of ERASE-multiple annotation enrichment (MAE) method for two SNP annotation data sources**

ERASE-MAE starts with finding the SNPs common (C) to the set of SNPs examined for ASE and the two input SNP annotation datasets. Each annotation dataset then follows the same but independent process: SNPs are randomly selected $10^4$ times from the non-ASEs to match the average read distribution of the ASEs in C. This randomization step generates z-scores from the corresponding SNP annotation dataset. The independent process ends at this step. The two z-scores for the observed set of ASEs (left and right branches of the flow diagram) are used to calculate the final z-score through a linear combination of both z-scores weighted by α and 1- α respectively, where α = a calibration parameter that represents the relative weight assigned to the SNP annotation dataset. The null distributions of random z-scores, on the other hand, are integrated as follows: randomly selected pairs of scores from both distributions are used to calculate a single random z-score identically as in the observed z-score (i.e. same α's used) so the mean and standard deviation can be calculated. A one right-tailed z-test is then used to obtain the enrichment p-value.

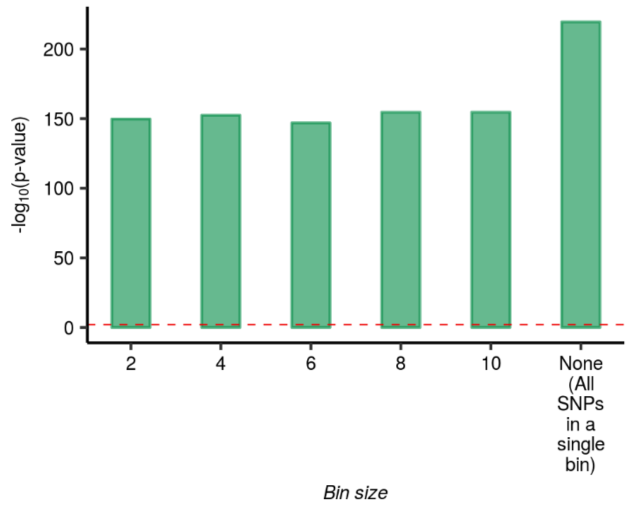


**Figure S2: Enrichment of eQTLs amongst ASEs using ERASE and the importance of examining enrichment in ASEs conditional on read depth.**

ERASE was run to check if it could detect the known enrichment of eQTLs in ASEs by applying it to gene-level eQTLs and ASEs derived from human post-mortem putamen tissue. Enrichment was observed when accounting for read depth, using the default bin size of 2. A stable p-value was observed when ERASE was run at increasing bin sizes of 4, 6, 8, 10. Not controlling for read depth (None) resulted in a highly inflated p-value. The red dashed line indicates the Bonferroni cut-off of 8.33 x $10^{-3}$.

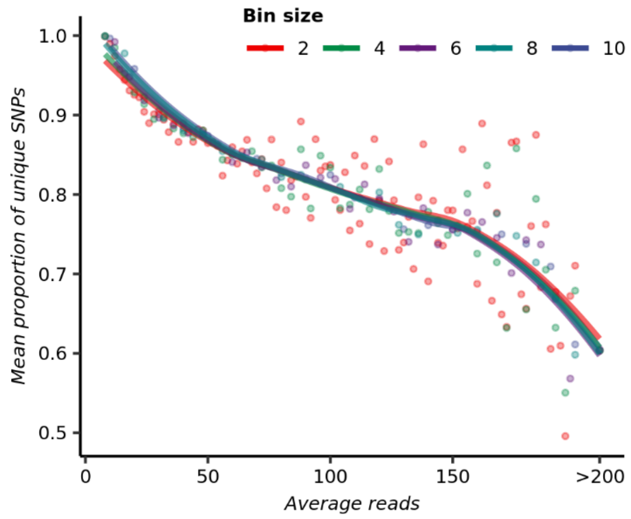


**Figure S3: Mean proportion of unique SNPs observed, running ERASE with varying bin sizes.**

ERASE was tested on gene-level eQTLs and ASEs derived from human post-mortem tissues at varying bin sizes. Each point on the plot represents the mean proportion of unique SNPs from those randomly selected for the corresponding average read depth, based on bin sizes of 2, 4, 6, 8, 10. The distribution of points was then approximated to a smooth curve with Loess regression. As expected, the proportion of unique SNPs chosen for the null population estimate decreases with an increase in the average read depth of the ASEs. However, bin sizes show small differences across the different values.

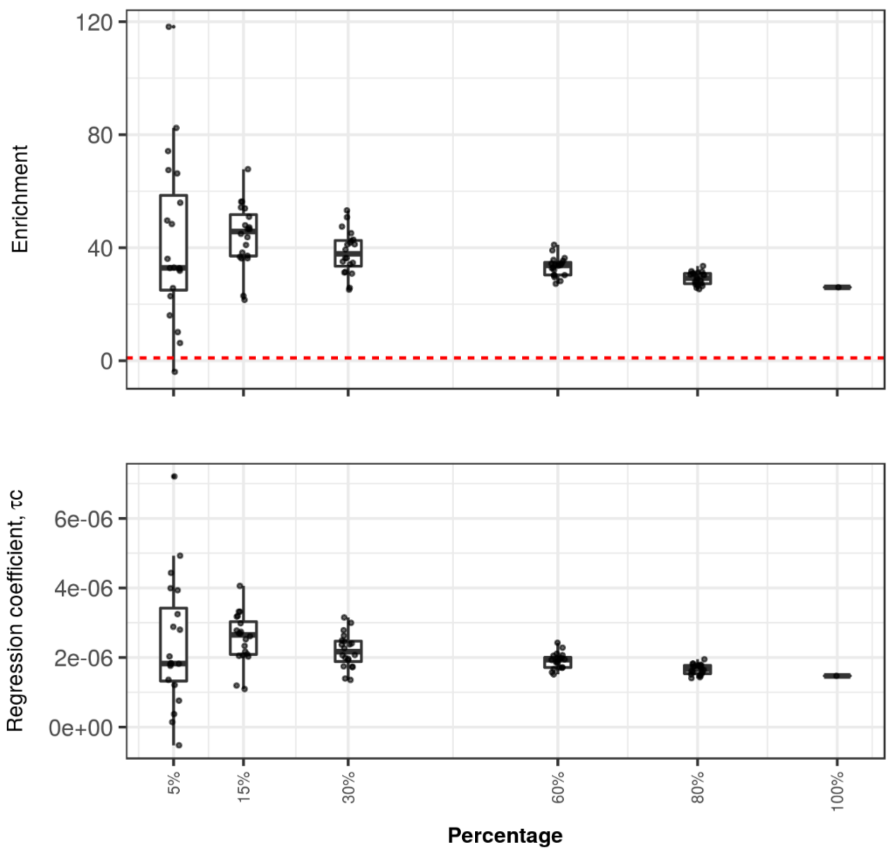


**Figure S4: Enrichment and regression coefficient plots with varying numbers of ASEs using stratified LDSC.** ASEs identified in the putamen and substantia nigra were repeatedly sub-sampled to a variety of sample sizes and stratified LDSC was applied across these groups. Enrichment is defined as the proportion of heritability explained by an annotation category divided by the proportion of SNPs within the category. The regression coefficient is a measure of how an annotation category contributes to trait heritability, conditional upon all the categories in the LDSC baseline model. Each box plot represents the enrichment or regression-coefficient values of 20 randomly sub-sampled datasets. The dashed red line indicates a value of 1 i.e. no enrichment. Corresponding co-efficient p-values are presented in Figure 3(b) and numerical results are reported in supplemental file S3.
